# Hitchhiking Microplastics Through Mosquitoes as Unseen Vehicles of Terrestrial and Aerial Transfer: Experimental Evidence

**DOI:** 10.64898/2026.06.01.729001

**Authors:** Mohammad Djaefar Moemenbellah-Fard, Mitra Boroomand, Shahin Saeedi, Zahra Derakhshan, Mehrzad Feilizadeh, Hamed Shouhanian, Sahar Souri Pilangorgi, Saied Nouri-Khorasani

## Abstract

**Background:** Microplastic (MP) pollution is an emerging environmental concern that extends beyond aquatic systems into terrestrial food webs. The mosquito *Anopheles stephensi*, a major malaria vector in Asia and the Middle East, undergoes complete metamorphosis and offers a useful model to examine cross-stage MP transfer. However, the physiological consequences of such exposure remain poorly understood.

**Methods:** Fluorescent polystyrene microplastics (1 µm) were administered at six concentrations (0–300 mg L^−1^) under controlled laboratory conditions. Larvae were reared on standard fish pellets, and adults were maintained on a 10% sucrose solution. Fluorescence microscopy tracked MP ingestion and transfer across life stages. Larval weight, developmental timing, and survival were analyzed using ANOVA and Chi-square tests.

**Results:** Ontogenic transfer of microplastics was confirmed from larvae to pupae (100%) and adults (94%). Increasing MP concentrations accelerated development by 1–2 days but significantly reduced survival, with mortality rising from 12% in controls to 64% at 300 mg L^−1^. Body weight declined consistently in a dose-dependent manner (p < 0.001).

**Conclusion:** This study provides the first experimental evidence of ontogenic MP transfer in *An. stephensi*. Microplastic exposure alters development and survival, potentially affecting mosquito population dynamics and vectorial capacity, highlighting the need to incorporate MP pollution into vector ecology models.

## 1. Introduction

Plastic pollution has become one of the most pervasive environmental challenges of the modern era, increasingly recognized for its persistence and capacity to impact both aquatic and terrestrial ecosystems. Alongside pharmaceuticals, organic dyes, and other anthropogenic polymer-based contaminants, plastic residues have contributed to a growing environmental burden that threatens the health of humans and other organisms[1, 2]. Among these pollutants, microplastics (MPs) as synthetic polymer particles smaller than 5 mm are of particular concern because of their stability, abundance, and bioavailability across ecological compartments[3]. These microscopic fragments are either generated through the gradual breakdown of larger plastics or intentionally produced for industrial use[4, 5]. Composed of polymers such as polyethylene, polyvinyl chloride, and polystyrene[6], MPs possess a high surface reactivity that allows them to adsorb persistent organic pollutants and pathogens, enhancing their ecological risk[7, 8].

Despite extensive documentation of MP contamination in marine and freshwater environments, little is known about their biological persistence within organisms that undergo complete metamorphosis. Recent research on *Culex* mosquitoes has shown that MPs ingested during larval feeding can persist through pupation and be detected in adults, indicating an ontogenic or cross stage transfer[9, 10]. This finding is ecologically significant because it suggests that aquatic larvae could act as biological vectors, transferring synthetic particles into terrestrial ecosystems via adult emergence. Moreover, it raises important questions about the ability of MPs to survive metamorphosis, a process involving extensive tissue reorganization, and the implications this persistence might have for the biological stability of plastic pollutants in living organisms[11].

*Anopheles stephensi*, the dominant malaria vector in Iran and parts of South Asia[12, 13], provides an ideal model to investigate this process. Its aquatic larvae frequently develop in water bodies contaminated with plastic waste, offering a natural interface between anthropogenic pollution and vector ecology. However, no study to date has traced the ontogenic transfer of MPs within *An. stephensi* Understanding whether microplastics can persist from larval to adult stages in this species is crucial not only for mapping pollutant pathways between ecosystems but also for exploring potential indirect effects on mosquito physiology and vectorial capacity.

Therefore, the present study aimed to trace and confirm the ontogenic transfer of fluorescent polystyrene microplastics from larvae to adults of *Anopheles stephensi* under controlled laboratory conditions. In addition to documenting the persistence of MPs across metamorphosis, secondary observations were made on developmental timing and survival to provide a broader ecological perspective on this phenomenon.

## 2. Materials and Methods

### 2.1. Mosquito rearing

The *An. stephensi* Shiraz strain, originally collected from the vicinity of Shiraz, Iran, was obtained from the insectarium colony maintained at School of Health, Shiraz University of Medical Science. The colony was reared under controlled laboratory conditions at 27 ± 1 °C, relative humidity of 70 ± 10%, and a 14:10 h light–dark photoperiod. Larvae were maintained in enamel trays containing dechlorinated tap water and were fed daily with commercial fish pellets (Tropical^®^, Iran) at a concentration of 1 g L^−1^. Adult mosquitoes were supplied *ad libitum* with a 10% sucrose solution. All experiments were conducted using healthy third instar larvae of uniform age (approximately 48 h post second molt).

### 2.2. Polystyrene microplastic microspheres

The procedure for microplastic preparation followed the method described by Simakova et al. with minor modifications[6]. Fluorescent green polystyrene microspheres (2.0 ± 0.2 µm diameter; density = 1.050 g cm^−3^; excitation/emission = 470/505 nm; Sigma-Aldrich, USA) were used in all experiments.

A stock suspension (2.5 mg mL^−1^) was prepared in 0.22 µm filtered distilled water and homogenized using a vortex mixer for 1 min to prevent particle aggregation. To ensure uniform dispersion, the stock suspension was washed thrice via centrifugation at 9000 rpm for 10 min, followed by resuspension in filtered distilled water after each cycle. All glassware was rinsed with filtered distilled water prior to use, and plastic materials were avoided to minimize secondary contamination.

### 2.3. Microplastics exposure design

Six treatment concentrations were prepared: 0 mg L^−1^ (control), 0.1 mg L^−1^, 1 mg L^−1^, 10 mg L^−1^, 100 mg L^−1^, and 300 mg L^−1^. Each treatment consisted of five replicates, each containing ten third instar larvae in 100 mL of purified water in a glass beaker. All groups received 100 mg of fish pellets during the exposure period.

To minimize potential airborne contamination, each beaker was covered with aluminum foil perforated with small holes to allow adequate aeration and gas exchange. The beakers were randomly positioned on the laboratory bench and gently rotated daily to ensure homogeneity of exposure. The experiment continued until adult emergence (approximately 10–12 days). The control group was maintained under identical conditions but without MPs.

### 2.4. Detection and quantification of MPs

Samples of fourth-instar larvae, pupae, and adults were collected from each treatment and rinsed twice with filtered distilled water to remove surface adhered particles. Specimens were then preserved in 70% ethanol in 1.5 mL Eppendorf tubes until analysis.

To assess the internal presence of MPs, individuals were digested using a wet peroxide oxidation (WPO) method modified from Lusher et al. [14]. Each specimen was treated with 35% hydrogen peroxide (H_2_O_2_) and 0.05 M FeSO_4_ (3:1 v/v) as a catalyst and incubated until complete dissolution of organic material was achieved.

Mature and pupae digestates were gently homogenized, and three 2 μL subsamples were randomly selected from each. Larval samples, due to their transparency, were directly mounted on glass slides and examined under an epifluorescence microscope (Lionheart FX, BioTek Instruments, USA) at 505/470 nm excitation/emission.

### 2.5. Statistical analyses

Prior to statistical analyses, data were tested for normality using the Shapiro–Wilk test and for homogeneity of variances using the Levene’s test. Normally distributed data with equal variances were analyzed by one-way ANOVA, followed by Bonferroni *post hoc* comparisons. Nonparametric data (*e.g*., survival counts) were evaluated using Chi-square tests. Statistical significance was set at *P* ≤ 0.05.

### 2.6. Quality assurance and contamination control

Comprehensive contamination control measures were implemented during all experimental and analytical steps. All glassware was cleaned with 0.22 µm filtered distilled water before use, and only glass or metal instruments were utilized. During mosquito rearing and exposure, all beakers were covered with perforated aluminum foil, allowing adequate ventilation while preventing airborne deposition of microplastics.

Sample preparation, digestion was conducted under a laminar flow hood. Laboratory personnel wore 100% cotton laboratory coats. No fluorescent MPs were observed in controls, confirming procedural integrity.

## 3. Result

### 3.1. Ontogenic transfer of microplastics

Fluorescence microscopy analysis revealed clear evidence of ontogenic transfer of polystyrene microplastics (MPs) throughout the developmental stages of *Anopheles stephensi* (Fig. 1). Fluorescence micrographs (Fig. 1a–c) confirmed ingestion and internal retention of green fluorescent particles in larval midguts and their subsequent translocation to pupal and adult tissues. The transfer rate from larvae to pupae reached 100%, while from pupae to adults it remained high at 94%. Control samples (Fig. 1d) were consistently free of detectable MPs, confirming the absence of background contamination and validating experimental sterility controls.

**Fig. 1.**
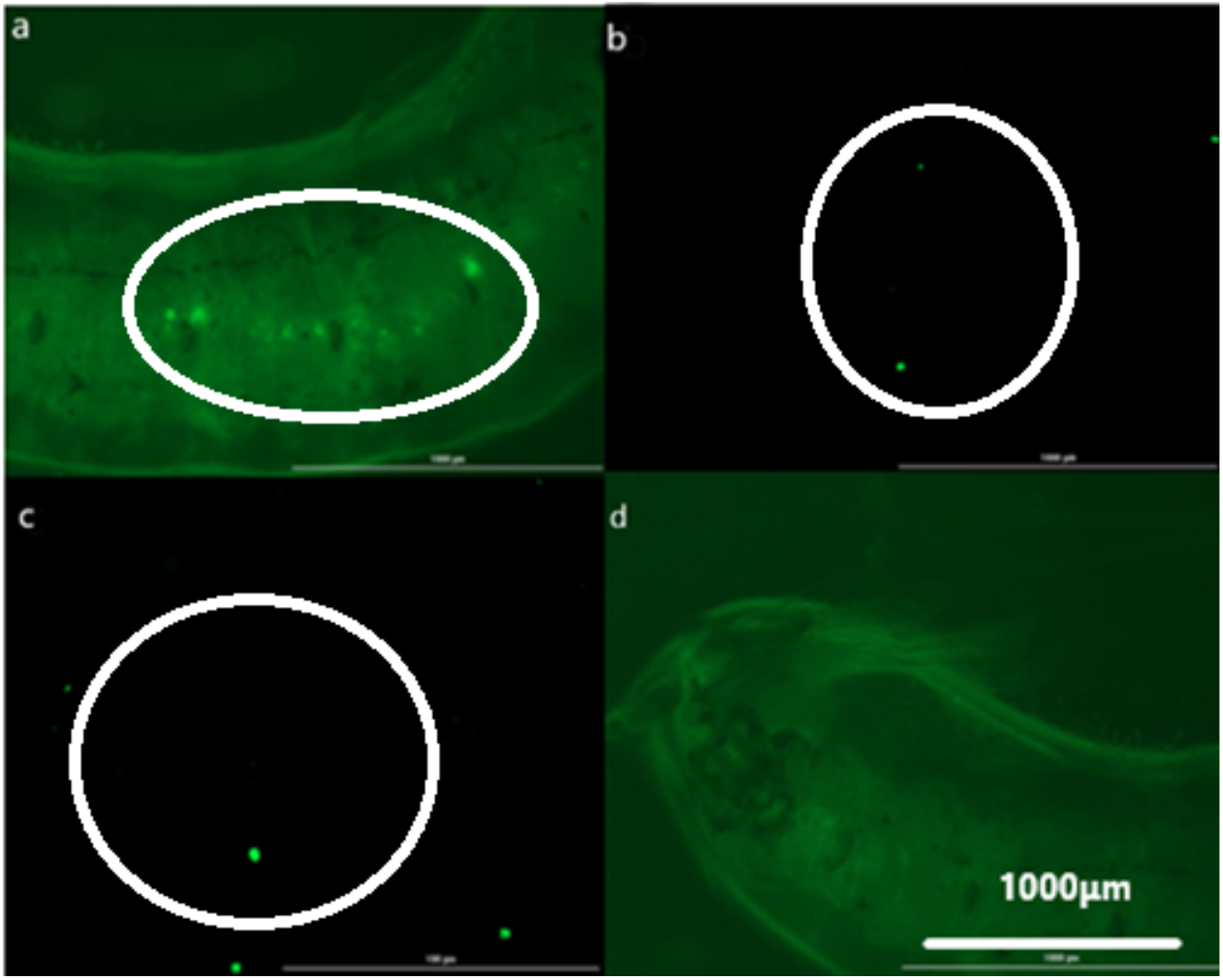
Microplastic particles found in (a) larvae (digestive system), (b) pupae, (c) adults, and (d) control. The scale bar is the same in all photos.

### 3.2. Rate of metamorphosis

Exposure to MPs significantly altered the developmental timing of *Anopheles stephensi* across larval and pupal stages. The transition from the third to fourth instar occurred earlier in exposed groups (Day 3–4; mean ≈ 4) compared with the control (Day 4–5; mean ≈ 4). Similarly, pupation was advanced by approximately 2 days, taking place on Day 6–7 (mean ≈ 6) in the experimental groups versus Day 8–9 (mean ≈ 8) in the control. Adult emergence (eclosion) was also accelerated, occurring between Days 9 and 10 (mean ≈ 9) among treated mosquitoes, whereas control adults emerged between Days 12 and 14 (mean = 12). One-way ANOVA confirmed a significant effect of MP concentration on total development duration (F (5, 54) = 12.84, *P* < 0.001), with post hoc comparisons showing that exposure levels ≥10 mg L^−1^ differed significantly from the control (*P* < 0.01).

### 3.3. Survival rate and body weight

Table 1 presents the survival and mortality data of *Anopheles stephensi* exposed to different concentrations of polystyrene microplastics. Mortality ranged from 12 % in the control group to 64 % at 300 mg L^−1^, with corresponding survival rates from 88 % to 36 %.

**Table 1.** Mortality and survival of Anopheles stephensi at different concentrations of polystyrene microplastics.

| Concentration<br>($\text{mgL}^{-1}$) | N* | Mortality<br>(Count) | Mortality (%) | Survival<br>(Count) | Survival (%) |
| --- | --- | --- | --- | --- | --- |
| Control | 50 | 6 | 12.0 | 44 | 88.0 |
| 0.1 | 50 | 23 | 46.0 | 27 | 54.0 |
| 1 | 50 | 26 | 52.0 | 24 | 48.0 |
| 10 | 50 | 19 | 38.0 | 31 | 62.0 |
| 100 | 50 | 27 | 54.0 | 23 | 46.0 |
| 300 | 50 | 32 | 64.0 | 18 | 36.0 |
| Total | 300 | 133 | 44.3 | 167 | 55.7 |

\*Each treatment contained 50 larvae (5 replicates × 10 larvae). Chi-square test: $\chi^2(5) = 32.97, P < 0.001$ .

Table 2 summarizes the mean body weight of mosquitoes across all experimental groups. The control group exhibited the highest mean weight (0.880 ± 0.258 mg), while the lowest mean weight (0.428 ± 0.149 mg) was recorded at 300 mg L^−1^. A gradual decline in body weight values was observed with increasing concentrations of microplastics (Figures 2).

**Table 2.** Descriptive statistics of *Anopheles stephensi* adult body weight (mg) following larval exposure to polystyrene microplastics.

| Concentration (mgL<sup>-1</sup>) | n | Mean (mg) | SD | SE | 95% CI (Lower-Upper) | Min | Max |
| --- | --- | --- | --- | --- | --- | --- | --- |
| Control | 50 | 0.88 | 0.258 | 0.037 | 0.808–0.952 | 0.6 | 1.2 |
| 0.1 | 50 | 0.67 | 0.144 | 0.02 | 0.631–0.709 | 0.4 | 0.9 |
| 1 | 50 | 0.612 | 0.139 | 0.02 | 0.572–0.652 | 0.4 | 0.9 |
| 10 | 50 | 0.582 | 0.135 | 0.019 | 0.543–0.621 | 0.3 | 0.9 |
| 100 | 50 | 0.5 | 0.153 | 0.022 | 0.457–0.543 | 0.2 | 0.9 |
| 300 | 50 | 0.428 | 0.149 | 0.021 | 0.386–0.470 | 0.2 | 0.8 |

One-way ANOVA: $F(5, 294) = 12.84$ , $P < 0.001$ . Each treatment contained 50 adults (5 replicates × 10 larvae).

**Fig. 2.**
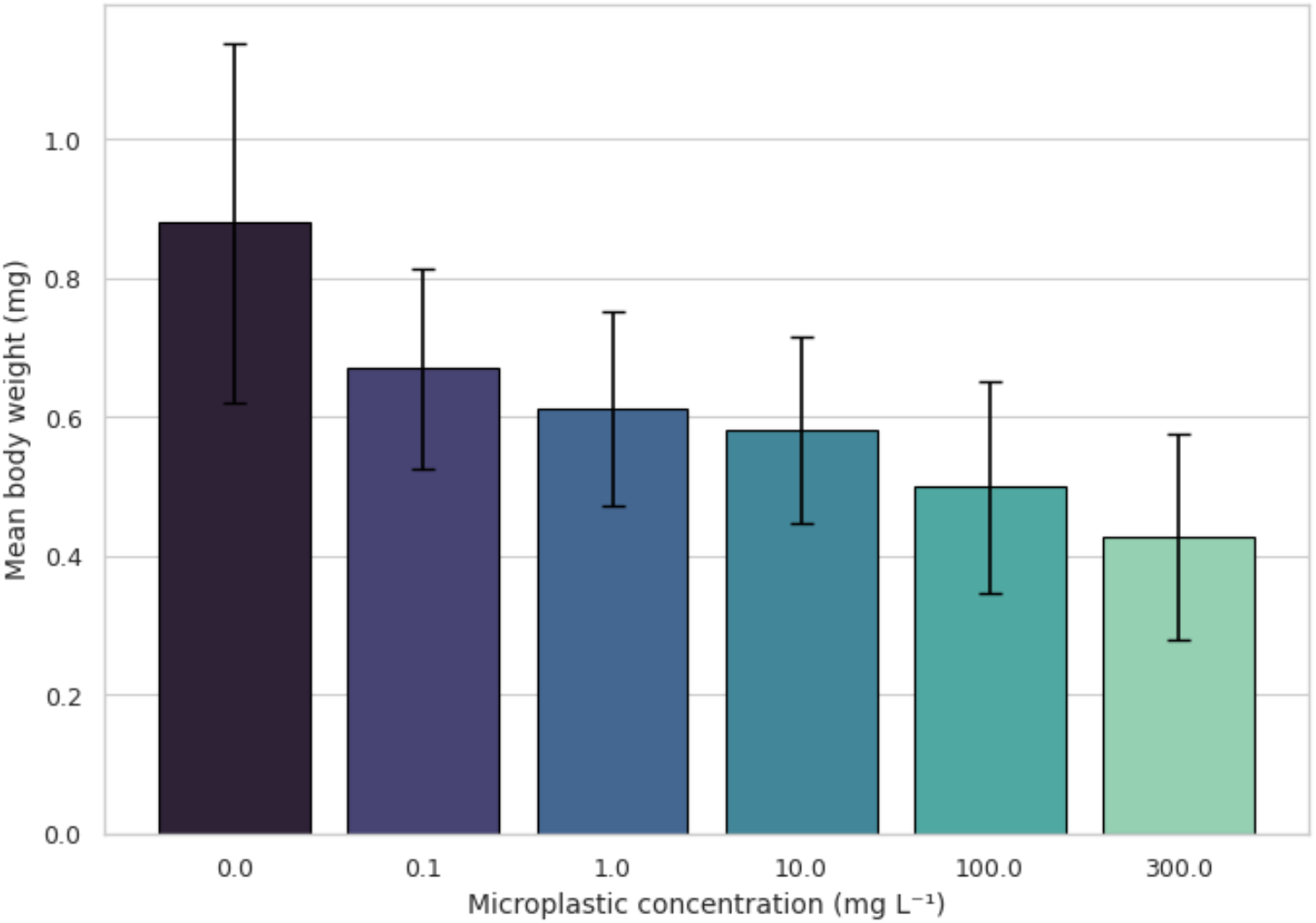
Mean (±SE) body weight (mg) of mosquitoes in the control and experimental groups. **0.0**: control

A statistically significant difference was detected in the mean body weight between the control and treatment groups (one-way ANOVA, *P*<0.001). Post hoc pairwise comparisons using the Bonferroni correction revealed significant differences in mean weight (W) between several dose levels. The control group (0 dose) showed significantly higher mean weight compared with all treated groups (0.1, 1, 10, 100, and 300 mgL^−1^; all *P*<0.05). No significant differences were observed among the lower dose groups (0.1, 1, and 10 mgL^−1^; *P*>0.05).

Higher doses (100 and 300 mgL^−1^) were associated with significantly lower mean weight compared with lower doses (0.1 and 1 mgL^−1^; *P* < 0.05). No significant difference was observed between the two highest dose levels (100 vs. 300 mgL^−1^; *P* > 0.05).

## 4 Discussion

The observed ontogenic transfer of MPs across larval, pupal, and adult stages of *Anopheles stephensi* highlights a biologically and ecologically significant route of plastic particle movement through aquatic-terrestrial boundaries. The stage dependent accumulation pattern, with maximal retention in fourth instar larvae and a gradual decline in subsequent stages, aligns with previous findings in *Aedes aegypti* and *Culex pipiens* that demonstrate similar life stage linked dynamics of microplastic uptake and retention[6, 9]. The 100% larval-to-pupal and 94% pupal-to-adult transfer efficiency observed here confirms that MPs are not merely transient contaminants but can persist through complete metamorphosis, thereby serving as a mechanism for aerial dispersal of aquatic pollutants[15]. From a mechanistic standpoint, the retention and translocation of MPs likely occur via gut associated tissues and hemolymph transfer during histolysis and histogenesis in metamorphosis, as reported in comparable studies on ontogenic MP transfer in mosquitoes and other holometabolous insects[16].

The significant reduction of MP load in adults compared to larvae suggests partial excretion or sequestration processes during pupation, consistent with findings in *Aedes aegypti*, where ecdysis and remodeling of gut epithelia resulted in partial particle clearance[17]. Ecologically, this transfer implies that mosquitoes act as biomechanical vectors for microplastics, linking aquatic and terrestrial ecosystems and potentially introducing MPs into food webs via predation on adult mosquitoes by insectivorous species [18].

The observed alteration in developmental timing of *Anopheles stephensi* under MPs exposure, although statistically significant, should be interpreted with caution. Given that the present study primarily aimed to document ontogenic transfer of MPs rather than explore physiological mechanisms, the earlier molting and eclosion events likely reflect indirect developmental modulation rather than a direct endocrine or molecular disruption. Similar non-mechanistic trends have been reported in *Aedes* and *Culex* species exposed to MPs, where subtle shifts in developmental timing were observed without definitive evidence of hormonal interference[19, 20]. The advancement of pupation and adult emergence by approximately 1–2 days at concentrations ≥10 mg L^−1^ suggests that MPs may impose a mild physiological or environmental stress, prompting accelerated development as a compensatory response, as previously proposed in other vector species [21]. However, without histological or endocrine data, it cannot be conclusively stated whether this response arises from altered hormone regulation (*e.g*., ecdysteroid or juvenile hormone activity), nutrient absorption changes, or stress induced metabolic acceleration [22]. Furthermore, while the developmental acceleration observed here may indicate stress induced plasticity, similar findings have not been consistent across mosquito taxa. For instance, Thormeyer & Tseng[20] reported no significant effects of environmentally realistic MP exposure on *Culex pipiens*, suggesting that responses may vary by species, polymer type, and experimental design.

Therefore, these observations should be viewed as preliminary evidence of potential developmental modulation under MP exposure rather than proof of a mechanistic causal link. From an ecological standpoint, earlier eclosion could theoretically influence mosquito population dynamics by altering aquatic terrestrial transition timing. Nonetheless, given the controlled laboratory context and the absence of field validation, any extrapolation to ecological or epidemiological impacts should remain speculative [18, 23]. Future work incorporating histological, hormonal, and transcriptomic assays would be valuable to confirm whether MPs directly influence the endocrine regulation of metamorphosis or merely induce stress related shifts in development pace.

The observed decline in both survival rate and body weight of *Anopheles stephensi* with increasing MP concentrations suggests a dose dependent physiological stress, though these findings should be interpreted cautiously given the observational scope of the present study. The reduction in survival from 88% (control) to 36% (at 300 mgL^−1^) is consistent with previous laboratory observations on *Culex* and *Aedes* mosquitoes exposed to polyethylene or polystyrene MPs, where higher concentrations induced moderate to severe mortality, likely through ingestion related stress and digestive disruption[9, 23]. One plausible explanation for the increased mortality and reduced weight is gut obstruction and impaired nutrient assimilation caused by physical accumulation of MPs in the alimentary canal.

Studies on *Aedes aegypti* and *Culex quinquefasciatus* have reported that polystyrene MPs accumulate in the midgut, altering digestion efficiency and leading to decreased biomass gain during larval development[24, 25]. However, without histological validation, it cannot be concluded that direct tissue damage or endocrine interference occurred in *An. stephensi* the observed effects may simply reflect energetic tradeoffs or feeding inefficiency under MP exposure. The progressive decline in body weight from 0.880 ± 0.258 mg in controls to 0.428 ± 0.149 mg at 300 mg L^−1^ mirrors findings from *An. gambiae* exposed to PET-based MPs, where larvae that survived to adulthood exhibited significantly lower mass and reduced fecundity[21]. Similar trends were also observed in *Culex quinquefasciatus* and *An. quadrimaculatus*, where ingestion of polystyrene spheres led to a measurable decrease in adult body weight and mating success[26]. From a toxicological perspective, the results support the hypothesis that MPs act as sublethal stressors rather than acute toxins. The statistically significant differences in body weight even at low concentrations indicate that chronic, low dose exposure may subtly impair growth performance. However, as Jones et al. and Thormeyer & Tseng [18, 20] emphasize, MP effects on mosquito physiology can vary depending on polymer type, particle size, and experimental realism.

## Conclusion

This study provides the first evidence of ontogenic transfer and sublethal physiological effects of MPs in *Anopheles stephensi*. Fluorescence microscopy confirmed the translocation of MPs across larval, pupal, and adult stages, demonstrating that microplastics can persist through complete metamorphosis. Developmental analyses revealed accelerated molting and eclosion in exposed groups, while higher MP concentrations significantly reduced survival and adult body weight. Together, these findings suggest that MPs act as chronic environmental stressors, altering developmental kinetics and energetic balance in mosquitoes. Although the present results are observational and lack histopathological confirmation, they underscore the need for mechanistic studies to elucidate how MPs influence insect physiology, vector ecology, and pollutant transfer across aquatic terrestrial interfaces.

## DECLARATIONS

### Data Availability Statement

The data supporting the results of this research are all contained within the manuscript.

### Ethical Approval

All authors consented to participate in this study, which was conducted based on the Declaration of Helsinki. The proposal was approved by the Ethics Committee of Shiraz University of Medical Sciences (IR.SUMS.SCHEANUT.REC.1402.099), and the research project code was 28467. This work was reviewed, approved, and funded to the main corresponding author (M.D.M-F) by Shiraz University of Medical Sciences (SUMS), Shiraz, Iran.

### Consent to Participate

Not applicable

### Consent to Publish

All authors have agreed to publish this manuscript.

### Credit Authorship Contribution Statement

S.S., M.B., Z.D., and M.D.M-F. conceptualized, and wrote the study design with a proposal. Z.D., M.F., and S.N-K. approved the proposal. S.S., M.B., S.N-K. and Z.D. facilitated the polystyrene purchase. They also carried out the mosquito collections, breeding, and experimental works. S.S.P. vindicated the statistical approaches. Expert validation and assistance on methods and data analysis was done by Z.D., M.F., H.S., and M.D.M-F., while S.S. and M.B. performed data collation, analyses, and draft preparation. M.D.M-F. approved the submitted version. All authors contributed to the final draft writing, review and approval of this manuscript.

## Acknowledgements

The authors thank the Vice-Chancellor for Research at Shiraz University of Medical Sciences (SUMS). This report was part of our Ph.D. students (Shahin Saeedi and Mitra Boroomand) ideas with a proposal number of 28467, and an ethical code of conduct (IR.SUMS.SCHEANUT.REC.1402.099) on behalf of these students awarded to their main supervisor (MD Moemenbellah-Fard.) of this research study. The authors are indebted to all our academic staff for their assistance in the fulfillment of this proposal at SUMS, Shiraz University, and Isfahan University of Technology, Iran.

